# A discovery platform to identify inducible synthetic circuitry from varied microbial sources

**DOI:** 10.1101/2023.10.13.562223

**Authors:** Clare M. Robinson, David Carreño, Tim Weber, Yangyumeng Chen, David T. Riglar

## Abstract

Gut microbes encode a variety of systems for molecular sensing and controlling conditional gene expression within the mammalian gut. Synthetic biology approaches such as whole-cell biosensing and ‘sense-and-respond’ therapeutics aim to tap into this vast sensing repertoire to drive clinical and pre-clinical applications. An ongoing constraint is the limited number of well-characterized inducible circuit components to specifically sense *in vivo* conditions of interest, such as disease. Here, we extend the flexibility and power of a biosensor screening platform using bacterial memory circuits encoded in a gut commensal *E. coli*. We construct libraries driven by potential sensory components derived from a combination of *E. coli* promoters or bacterial two-component systems (TCSs) sourced from diverse gut bacteria. Each is tagged with unique DNA barcodes using a pooled construction method. Using our pipeline, we evaluate sensor activity and performance heterogeneity across *in vitro* and *in vivo* conditions including using a mouse inflammation model. We demonstrate the method’s ability to identify biosensors of interest, including the identification of unannotated TCSs. Following the optimisation of library construction, analysis, and delivery to account for the challenges of working with engineered bacteria within a conventional mammalian gut microbiome, we identify and validate several further biosensors of interest responding to the murine gut environment, and specifically during inflammatory conditions. This approach can be applied to transcriptionally activated sensing elements of any type and will allow for rapid development of new biosensors that can advance synthetic biology approaches for complex environments.

## Introduction

A longstanding vision for synthetic biology is the development of bacterial whole-cell biosensors and ‘sense-and-respond’ live biotherapeutic products. These systems promise to autonomously sense and induce reporters or therapeutics under defined spatial, dietary and disease conditions ^1,2^. Potential applications include diagnostics, disease monitoring and biotherapeutics ^2^. However, well-characterised biosensing circuits for the mammalian gut remain limited, hindering the complete fulfilment of this vision^3,4^.

Synthetic genetic memory circuits, which convert transient signals into sustained responses, for example through transcriptional switches ^5,6^ or DNA editing ^7-10^, are powerful tools for non-invasive biosensor reporting and actuation. In rodent models, such circuits have been used to successfully track inflammatory signals ^6,8,10,11^, record microbial response to dietary perturbations ^7,8^, detect tumour DNA ^12^, and actuate sustained biotherapeutic secretion ^10^. However, due to the extended, often permanent, nature of the response, input signals typically need to show low background activation combined with strong induction dynamics for best compatibility with these approaches.

We previously developed a high-throughput memory system (HTMS) in mouse commensal *Escherichia coli* NGF-1 ^13^ to screen *E. coli* promoters as biosensor candidates within the mouse gut ^11^. We now expand the power and flexibility of this screening platform to accommodate a diversity of sensory components from varied microbial sources (Figure 1A). These include multi-gene heterologous two-component systems (TCSs), for which we develop a custom computational pipeline for rapid identification of cloning-compatible systems, and shorter native promoter-only-based sensors, identified from previous studies to sense conditions of interest ^8,11^. Sensors are constructed by multiplexed golden gate assembly and integrated into the HTMS chassis (Figure 1B). Incorporating unique DNA barcodes into each sensor allows simultaneous testing of variable-sized sensors and flexibility for library combination and recombination.

**Figure 1.**
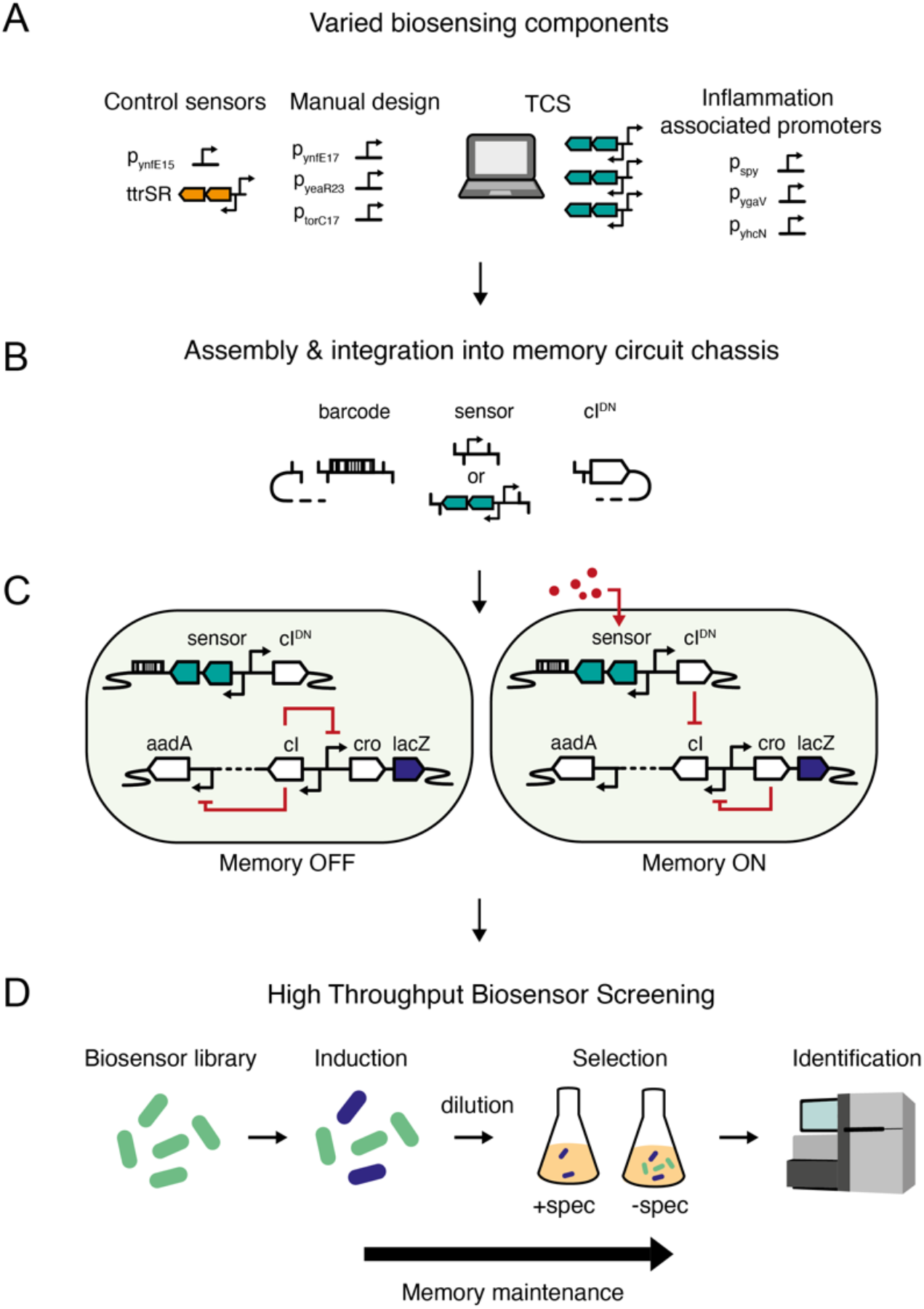
A high-throughput transcriptional screening pipeline based on an engineered memory circuit to identify novel inducible circuits. A) Various sensing components can be combined and B) assembled, along with unique DNA barcodes, by multiplexed golden gate assembly to drive switching of C) a *λ*-phage-based memory circuit. In the presence of an activating stimulus, cI^DN^ is expressed and inhibits *cI*, de-repressing *cro* and allowing expression of reporter genes *lacZ* and *aadA*. Cro expression maintains this memory ON state irrespective of ongoing exposure to the activating stimulus. D) A library of memory-linked bacterial biosensors can be rapidly screened for activated sensors by growth in spectinomycin selective media followed by sequencing of recovered gDNA for identification.

Memory recording is based on *λ*-phage’s lysis-lysogeny switch ^5^. In the chassis’ memory ‘OFF’ state, *cI* represses β-galactosidase (*lacZ*) and streptomycin/spectinomycin adenyltransferase (*aadA*) reporter cassettes (Figure 1C, left). Upon exposure to an inducing signal, the sensing component triggers expression of cI dominant negative (cI^DN^), switching cells into memory ‘ON’ states, in which the repression of *cro, lacZ* and *aadA* is relieved (Figure 1C, right). Repression of *cI* by cro maintains switching status irrespective of the presence of the induction signal, resulting in cellular ‘memory’. Activated sensors can be identified from a large library through selection by growth in spectinomycin followed by massive parallel sequencing (Figure 1D).

Bacterial TCSs are modular signalling pathways capable of detecting various molecular signals ^14^. They are typically composed of a membrane-bound histidine kinase (HK) and partner response regulator (RR) that control the transcription of target genes ^15^. While it is not expected that all TCSs will function in a heterologous host exactly as they do in their native host, we and others have successfully transferred sensing capacities between bacterial species. We previously developed a memory biosensor to the inflammatory biomarker tetrathionate, driven by a heterologously expressed TCS from *Salmonella enterica* subsp. *enterica* serovar Typhimurium (herein *S*. Typhimurium), which controlled *λ*-memory induction through the diverging promoter for the neighbouring ttr_BCA_ operon ^6^. Given the ability of TCSs to facilitate non-native sensing capacities in heterologous hosts, they are ideal targets for sensors to diversify engineered chassis biosensing capacity using a screening approach.

Here, we build two new libraries of sensor components: the first focused on TCSs from diverse bacterial origins (Library 1) and a second predominantly containing *E. coli* derived sensors targeted to be responsive to the mammalian gut (Library 2). We test these libraries *in vitro* and *in vivo* in the murine gut, identifying and validating several new sensors of interest. Included are those responsive to the gut environment both generally and specifically during inflammation. Together, this work advances methodology for the application of library-based screens of synthetic biology components in *in vivo* gut contexts and validates the power of this approach for identifying context-dependent sensors that can actuate synthetic circuits within the mammalian gut.

## Results

### Construction of bacterial biosensor libraries encoding diverse native and heterologous sensing machinery

Most HKs are encoded within 200bp of their partner RR ^16^. Where a HK-RR pair sit next to a diverging gene, we term this a ‘grouped’ TCS (Figure 2A). Many grouped TCSs regulate their adjacent and divergent genes. Of a total of 30 known TCS pairs in *E. coli* MG1655, 13 lie in this ‘grouped’ formation, and of these, 10/13 regulate the divergent, adjacent gene based on Ecocyc annotations ^17^. We, therefore, focused on these ‘grouped’ TCSs for biosensor library expansion due to the ability to replicate complete sensing machinery (HK, RR and putatively regulated promoter sequence) in a single PCR for transfer to the heterologous *E. coli*.

**Figure 2:**
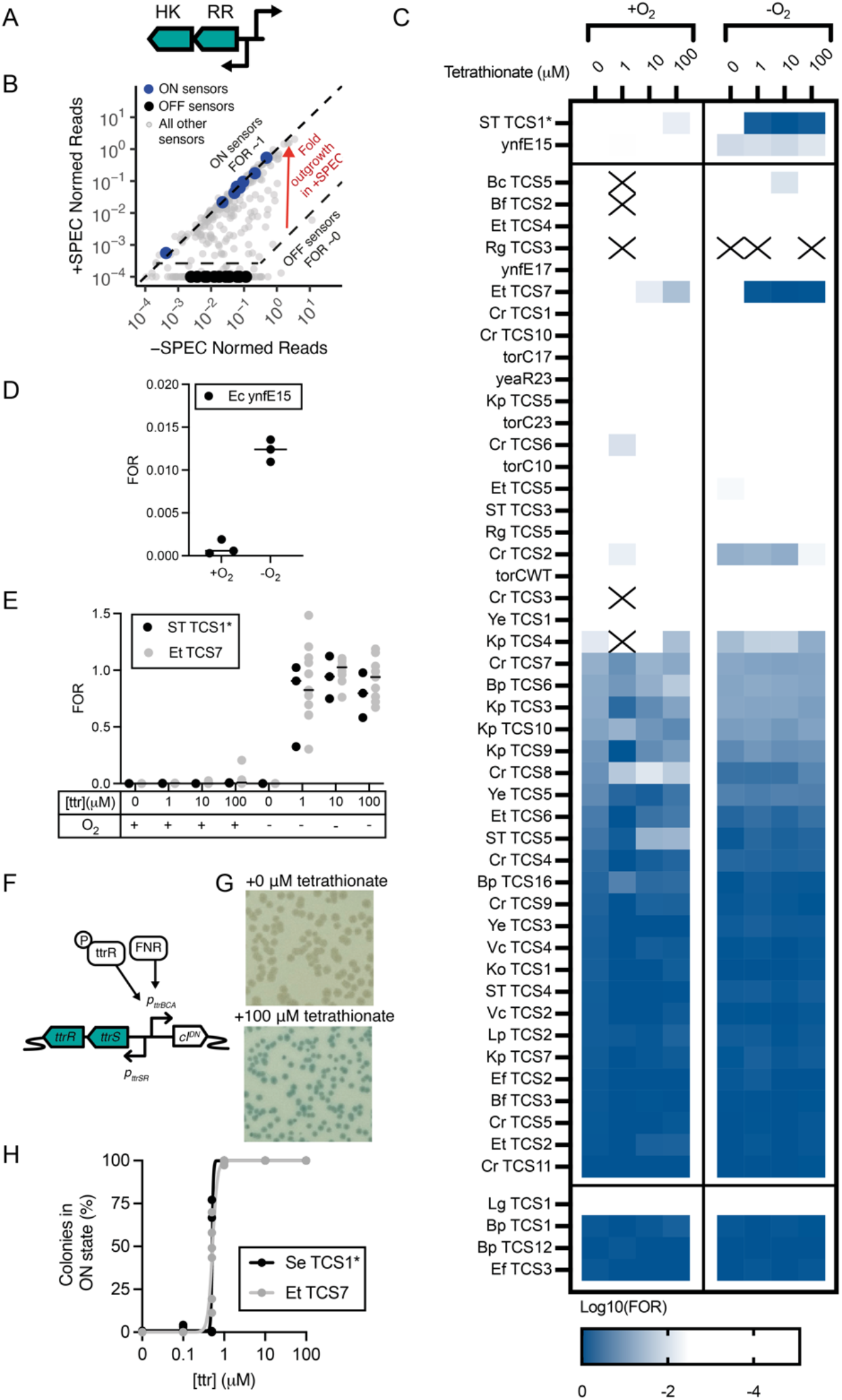
Barcoded biosensor screening of Library 1 identifies specific biosensors *in vitro*. A) ‘Grouped’ TCS arrangements are ideal for straightforward cloning of libraries encoding heterologous sensing components together with a probable regulated promoter to trigger memory formation. B) Normalised reads of a typical sample, with ON and OFF control sensors highlighted. ON sensors will have equal growth in the presence (+SPEC) or absence (-SPEC) of spectinomycin selective media, resulting in a fractional odds ratio (FOR) of ∼1. OFF sensors will not grow in spectinomycin selective media (+SPEC), resulting in a FOR of ∼0. C) Average log10 transformed fractional odds ratio of all sensors within the library in the presence and absence of oxygen and varying levels of tetrathionate. D) Ec ynfE15 sensors showed increased activation under anaerobic conditions. E) Sensor activation of barcodes from Et TCS7, ST TCS1* under anaerobic conditions across all oxygen and tetrathionate combinations tested. For D) and E) each point represents an individual barcode’s FOR, with medians marked. F) Putative sensor regulation of both Et TCS7 and ST TCS1* G) Et TCS7 was individually cloned (DTR302) and tested by exposure to anaerobic conditions in the presence and absence of 100μM tetrathionate and indicator plating, confirming the sensor is responsive to tetrathionate under anaerobic conditions. H) Response dynamics across various tetrathionate concentrations were similar to memory driven by ST TCS1*(DTR277).

To enable rapid library design, we developed a computational pipeline to identify grouped TCSs from available genomic sequencing data (see methods). Using this pipeline directed against 17 bacterial species for which we had access to both full genome sequences and gDNA of exact or closely related strains, we identified 153 ‘grouped TCSs’, 146 of which were compatible with our downstream cloning strategy (Table 1). Identified TCSs were cloned so that the divergent promoter would drive cI^DN^ expression (Figure 1B). Pooled golden gate assembly tagged each sensor with a 106bp randomised nucleotide hypervariable barcode flanked by sequencing adaptor sites for unique identification. The plasmid library was then transformed, and genome integrated into *E. coli* NGF-1 encoding the high-throughput memory system (PAS811) ^11^ (Supplementary Figure 1A).

**Table 1:**
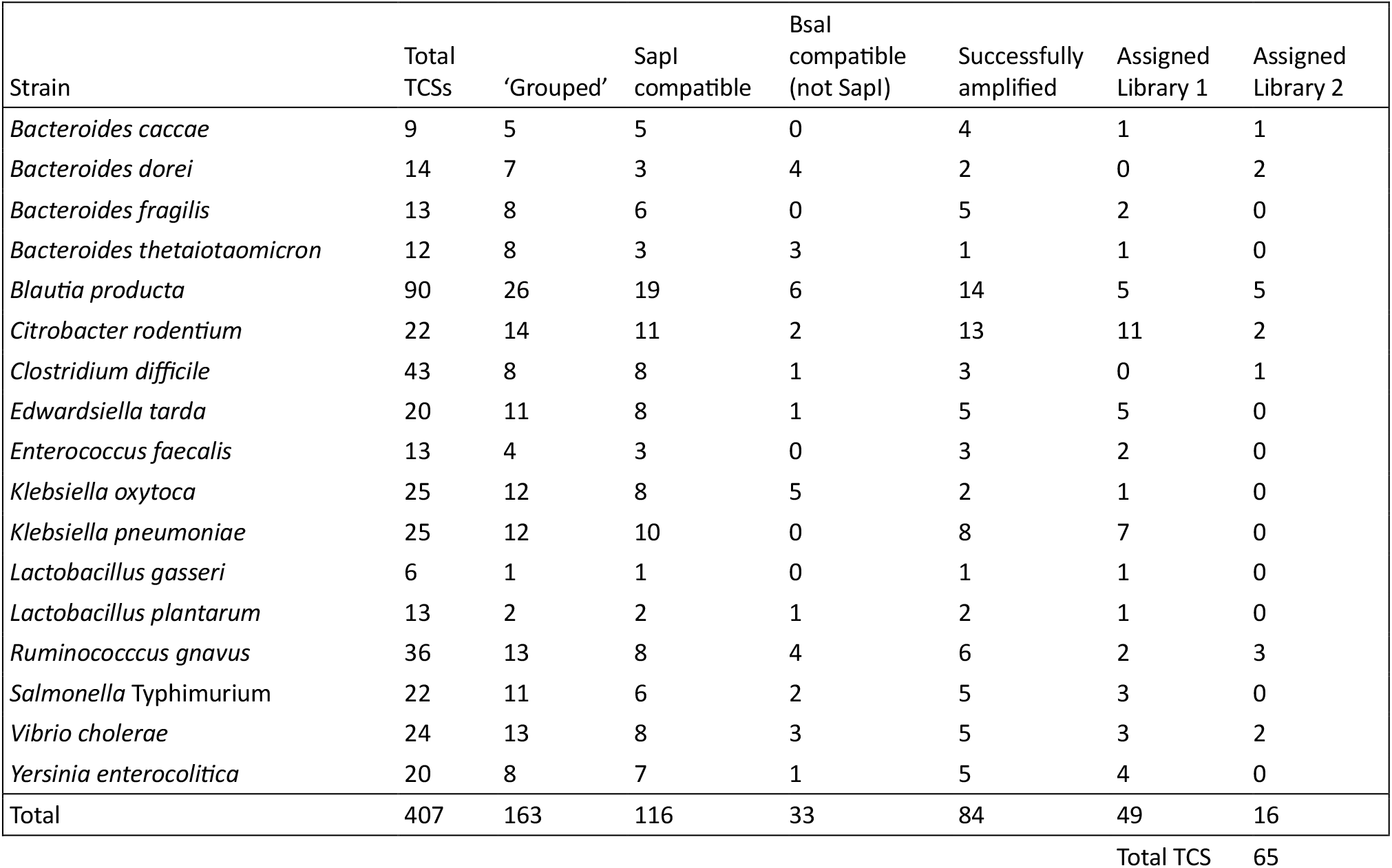
Compatible TCSs computationally identified and cloned from diverse gut bacteria.

| Strain | Total TCSs | 'Grouped' | SapI compatible | BsaI compatible (not SapI) | Successfully amplified | Assigned Library 1 | Assigned Library 2 |
| --- | --- | --- | --- | --- | --- | --- | --- |
| *Bacteroides caccae* | 9 | 5 | 5 | 0 | 4 | 1 | 1 |
| *Bacteroides dorei* | 14 | 7 | 3 | 4 | 2 | 0 | 2 |
| *Bacteroides fragilis* | 13 | 8 | 6 | 0 | 5 | 2 | 0 |
| *Bacteroides thetaiotaomicron* | 12 | 8 | 3 | 3 | 1 | 1 | 0 |
| *Blautia producta* | 90 | 26 | 19 | 6 | 14 | 5 | 5 |
| *Citrobacter rodentium* | 22 | 14 | 11 | 2 | 13 | 11 | 2 |
| *Clostridium difficile* | 43 | 8 | 8 | 1 | 3 | 0 | 1 |
| *Edwardsiella tarda* | 20 | 11 | 8 | 1 | 5 | 5 | 0 |
| *Enterococcus faecalis* | 13 | 4 | 3 | 0 | 3 | 2 | 0 |
| *Klebsiella oxytoca* | 25 | 12 | 8 | 5 | 2 | 1 | 0 |
| *Klebsiella pneumoniae* | 25 | 12 | 10 | 0 | 8 | 7 | 0 |
| *Lactobacillus gasseri* | 6 | 1 | 1 | 0 | 1 | 1 | 0 |
| *Lactobacillus plantarum* | 13 | 2 | 2 | 1 | 2 | 1 | 0 |
| *Ruminococcus gnavus* | 36 | 13 | 8 | 4 | 6 | 2 | 3 |
| *Salmonella* Typhimurium | 22 | 11 | 6 | 2 | 5 | 3 | 0 |
| *Vibrio cholerae* | 24 | 13 | 8 | 3 | 5 | 3 | 2 |
| *Yersinia enterocolitica* | 20 | 8 | 7 | 1 | 5 | 4 | 0 |
| Total | 407 | 163 | 116 | 33 | 84 | 49 | 16 |

Total TCS 65

Barcodes were identified and associated with their corresponding sensing components by a combination of massive parallel long- and short-read sequencing (Supplementary Figure 2A). Sequencing confirmed successful barcode assignment for 49 unique SapI compatible TCSs in the first instance. We supplemented these strains with 9 additional individually cloned ‘curated’ sensors identified in our previous studies ^6,11^ to afford ‘Library 1’ (Table 1, Data File 1). Curated sensors included two controls: i) a *S*. Typhimurium *ttrSR-*P_*ttrBCA*_ sensor, edited with a synthetic ribosome binding site, (ST TCS1*) that previously showed response to the inflammatory biomarker tetrathionate ^6^, and ii) the *E. coli* Nissle 1917 P_*ynfEFGH*_ promoter edited with a synthetic 5’ untranslated region and ribosome binding site, MCD15 (Ec ynfE15)^18^ that previously showed switching under *in vitro* anaerobic growth conditions and in the murine gut ^11^. Library 1 contained 504 assigned unique barcodes linked to these 58 sensors.

Building on the flexibility of the method to easily combine sensors, barcodes and libraries, we subsequently cloned a set of additional triggers, targeting those responsive to gut and gut inflammation related signals. These included *E. coli* promoter regions previously induced in mouse models, including during inflammation ^8^; 2) computationally identified TCSs that were not assigned in Library 1 construction, including those with BsaI but not SapI compatibility; and 3) several additional curated sensors of interest (Table 1, Data File 1). Library 2 contained 10,063 assigned unique barcodes linked to 17 additional TCSs and 72 *E. coli* promoter regions of interest.

The number of assigned barcodes per sensor ranged between 1 and 44 (median 5) for Library 1 and 1 and 1001 (median 52) for Library 2 (Supplementary Figure 2B). Each library demonstrated similar abundance profiles (Supplementary Figure 2C). Between the two libraries we confirmed at least 1 barcode for ∼80% of successfully PCR-amplified TCS sensor regions with this method (Table 1).

### In vitro validation confirms barcoded library biosensor screening capabilities

To quantify the state of each sensor we compared barcode sequencing read counts for each barcode from samples split and grown in the presence or absence of spectinomycin (Figure 2B). Only ON sensors grow with spectinomycin due to *aadA* expression (Figure 1C-D). Degree of switching is represented as a fractional odds ratio (FOR) by normalisation to reads from known positive and negative strains (see methods) (Figure 2B). FOR ∼1 corresponds to an ON sensor, and ∼ 0 to an OFF sensor.

To validate barcoded library screening, we cultured pooled sensors from Library 1 *in vitro* with varying tetrathionate concentrations and with or without oxygen (Figure 2C). Various sensors showed differential activation between the conditions. As expected, around half of the sensors were consistently ON in all conditions, suggesting a level of expression from these promoters during *in vitro* growth, resulting in activation of the memory circuit under all conditions. Mirroring our previous results, Ec ynfE15 sensors had elevated odds ratios in anaerobic conditions (Figure 2C-D) ^11^ and ST TCS1* sensors activated with tetrathionate in the absence of oxygen (Figure 2C) ^6^. Together, these results demonstrate the functionality of the barcoding approach to combine sensors of diverse size and origins.

### Barcode screening successfully identifies unannotated biosensor candidates

The 22 barcodes linked to *Edwardsiella tarda* TCS7 (Et TCS7) were also consistently induced at levels comparable to ST TCS1* across the conditions tested (Figure 2C, E). While the available gene annotations for the strain used for TCS extraction (*E. tarda* KC-Pc-HB1) did not predict this behaviour, BLAST searches identified homology to a putative tetrathionate reductase promoter region with its associated TCS in *E. tarda* strains Et54^19^ and FL6-60 ^20^ (Figure 2F). Et TCS7 was therefore cloned as an individual strain (DTR302) and its response to tetrathionate was confirmed by plating of induced cultures on X-gal indicator plates for blue-white colony screening. When grown anaerobically, both DTR302 and an individual strain triggered by ST TCS1* (DTR277) demonstrated strong induction in the presence of >1μM tetrathionate (Figure 2G-H). This demonstrates the ability of our screening pipeline to identify unannotated biosensors with specific sensing functions using our screening approach.

### Barcode and strain diversity is maintained over short periods within the murine gut

We have previously demonstrated robust *E. coli* NGF-1 colonisation of the conventional mouse gut, typically following a dose of streptomycin to reduce colonisation resistance, but in select cases also without pre-treatment ^5,6^. To test colonisation characteristics in our current facility, we administered sensor Library 1 by oral gavage (∼10^11^ bacteria/mouse) to groups of 2 healthy, conventional mice 24 hours following administration of streptomycin antibiotics (1mg or 5mg/mouse) or mock treatment (Supplementary Figure 3A). Faecal bacterial CFU counts reflected streptomycin provision, with engineered bacteria in untreated animals dropping below the detection limit at day 3, and in mice administered 1mg streptomycin on day 4-5 (Supplementary Figure 3B). Engineered *E. coli* successfully colonised mice provided with 5mg streptomycin throughout the full 53-day experiment (Supplementary Figure 3B).

Strain diversity, as estimated by percentage of barcodes identified via sequencing, was high in all animals at day 1 post-administration but not measurable by day 4 in untreated and 1mg streptomycin-treated mice as no engineered bacteria were recovered from these animals on these days (Supplementary Figure 3C). Even in 5mg streptomycin-treated mice that were effectively colonised, barcode and sensor diversity dropped quickly following administration (Supplementary Figure 3C and D). By day 4 only 3 – 4 % of total barcoded strains remained, with a handful of strains dominating colonisation within each mouse (Supplementary Figure 3E). Such colonisation bottlenecks are consistent with other work examining barcoded isogenic strains in the mouse gut ^21,22^. Consequently, all further library analyses focussed on short (2-day) experimental timeframes to maintain barcode diversity.

### Barcode screening successfully identifies biosensors in vivo

To test sensor activation in the gut, we administered Library 1 sensors by oral gavage (∼10^12^ bacteria/mouse) to mice (n = 4) 24 hours following administration of 5mg streptomycin (Figure 3A). Nearly all (>98%) barcodes and sensors (100%) were recovered on both days (Figure 3B-C). Various sensors showed elevated switching in animals compared to gavage cultures (Figure 3D; full data set: Supplementary Figure 4A). Consistent with previous testing ^11^, Ec ynfE15 strains showed odds ratio increases most prominently by day 2 (Figure 3E). The top-ranked activated sensor was *Citrobacter rodentium* TCS7 (Figure 3F), identified as the *ygiW* gene promoter paired with the adjacent and opposing TCS, QseCB. Expression of *ygiW* is regulated by the QseB homologue, PreA, in *S*. Typhimurium ^23^, and deletion of the HK QseC in K. pneumonia resulted increase of *ygiW* expression, suggesting the gene is directly or indirectly regulated by the QseBC system in this species also ^24^. However, in *E. coli*, expression of *ygiW* has not been linked to QseB, but rather hydrogen peroxide and cadmium stress ^25^, the gluconate utilisation response regulator, GlaR, and the leucine responsive global transcriptional regulator, Lrp (Figure 3G) ^17^.

**Figure 3:**
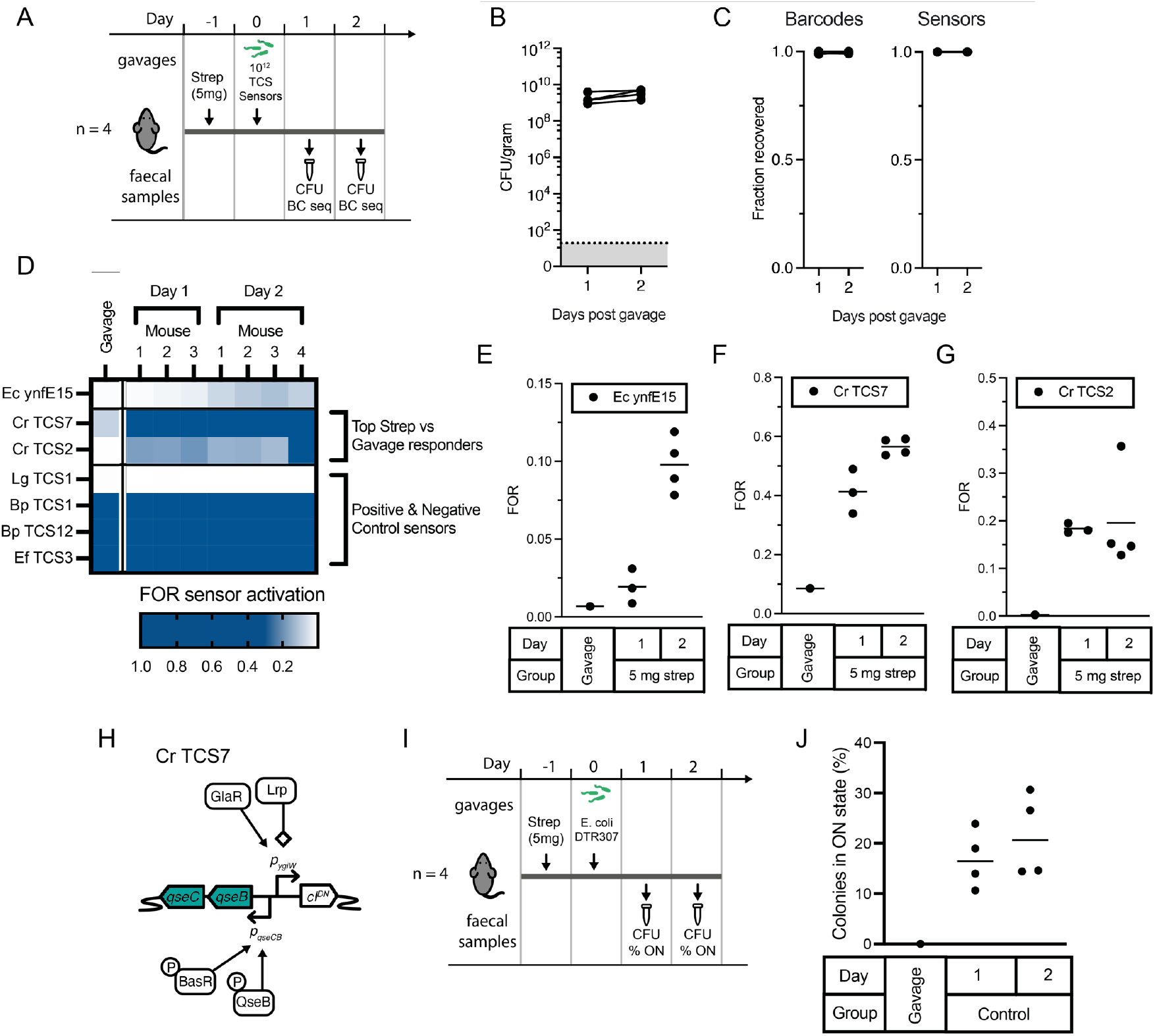
Barcoded memory screening of Library 1 successfully identifies biosensors for in vivo growth. A) Library 1 was administered to mice 1 day post-administration of 5 mg streptomycin (n=4). Faecal samples were collected for colonisation and memory switching analyses the two days following. B) Colonisation levels and C) fractions of recovered barcodes and sensors showed high retention in all mice. D) Sensor activation of top-ranked in vivo responsive sensors, and control sensors in mice on days 1 and 2 post library administration. Full dataset is shown in Supplementary Figure 4. E) The response of the control sensor, Ec ynfE15 and top-ranked in vivo responsive sensors F) Cr TCS7 and G) Cr TCS2 to the gut environment. Panels show FOR of faecal pellets from each mouse & gavage samples with mean response marked. H) Potential regulatory control of Cr TCS7 (QseBC-P_ygiW_) based on regulation of C. rodentium and E. coli homologues. I) Experiment timeline for testing the response of Cr TCS7 (DTR307) within the mouse gut. J) Percentage of activated sensor colonies from plating of faecal contents on days 1 and 2 after sensor administration, panel shows counts from plated individual mouse faecal pellets and the gavage sample.

To validate the observed sensor response, we cloned Cr TCS7 as an individual strain (DTR307) and delivered it to mice via oral gavage 24 hours following streptomycin treatment (5mg) (Figure 3H). DTR307 response to the gut was tested over the 48 hours following administration via plating of faecal samples on indicator plates (Figure 3I). Similarly to our library-based screen, the sensor was off in the gavage sample, and showed consistent switching in ∼20% of bacteria collected from faecal pellets from all mice on both days post gavage (Figure 3J). This validates the signal seen within the library-based screen and demonstrates the ability for the screening pipeline to identify sensors of interest within the mouse gut.

Another biosensor of interest ranked as showing higher response within the murine gut than our control Ec ynfE15 is Cr TCS2, identified as the *C. rodentium* intergenic region leading to *citC* with adjacent TCS DpiAB (Figure 3D, G, Supplementary Figure 5A-C). We further observed Ye TCS1, the *Y. enterocolitica* intergenic region leading to *macA* with adjacent TCS CpxAR (Supplementary Figure 4D-F), showing response to the gut, albeit below the threshold of the positive control Ec ynfE15. Homologous TCSs to these systems in *E. coli* are well characterised, with known inducers. When cloned and tested *in vitro* as individual sensors we confirmed the response of these heterologous sensors to known regulating signals for their cognate *E. coli* homologues (Supplementary Figure 5A-B, D-F). For Cr TCS2 we additionally tested truncated variants of the sensor, lacking the heterologous *C. rodentium dpiA* and *dpiB* genes, or DpiA binding sites (Supplementary Figure 5A-C). We observed a response to citrate in the truncated mutant lacking *dpiA* and *dpiB*, suggesting native *E. coli* host machinery is capable of regulation of the non-native intergenic promoter region, but no response to citrate upon the removal of DpiA binding sites, confirming regulation by DpiA is necessary for citrate response (Supplementary Figure 5C).

### Identification and validation of inflammation responsive biosensors

To specifically target inflammation-responsive circuit development, we combined Library 1 with Library 2 – the latter being predominantly built using *E. coli* promoters previously associated with response in the murine gut including during inflammation ^8^. We administered the combined libraries via oral gavage to mice (n=4 per group; ∼10^11^ bacteria/ mouse) pre-treated with streptomycin 24 hours prior (5mg/mouse) and with or without 3% DSS in their drinking water for the preceding 4 days to induce intestinal inflammation (Figure 4A). Colonisation levels were high in all animals and similar across both groups (Figure 4B). Reduction of colon length at endpoint and loss of body weight throughout the experiment confirmed the expected impact of DSS in treated animals (Figure 4C-D).

**Figure 4:**
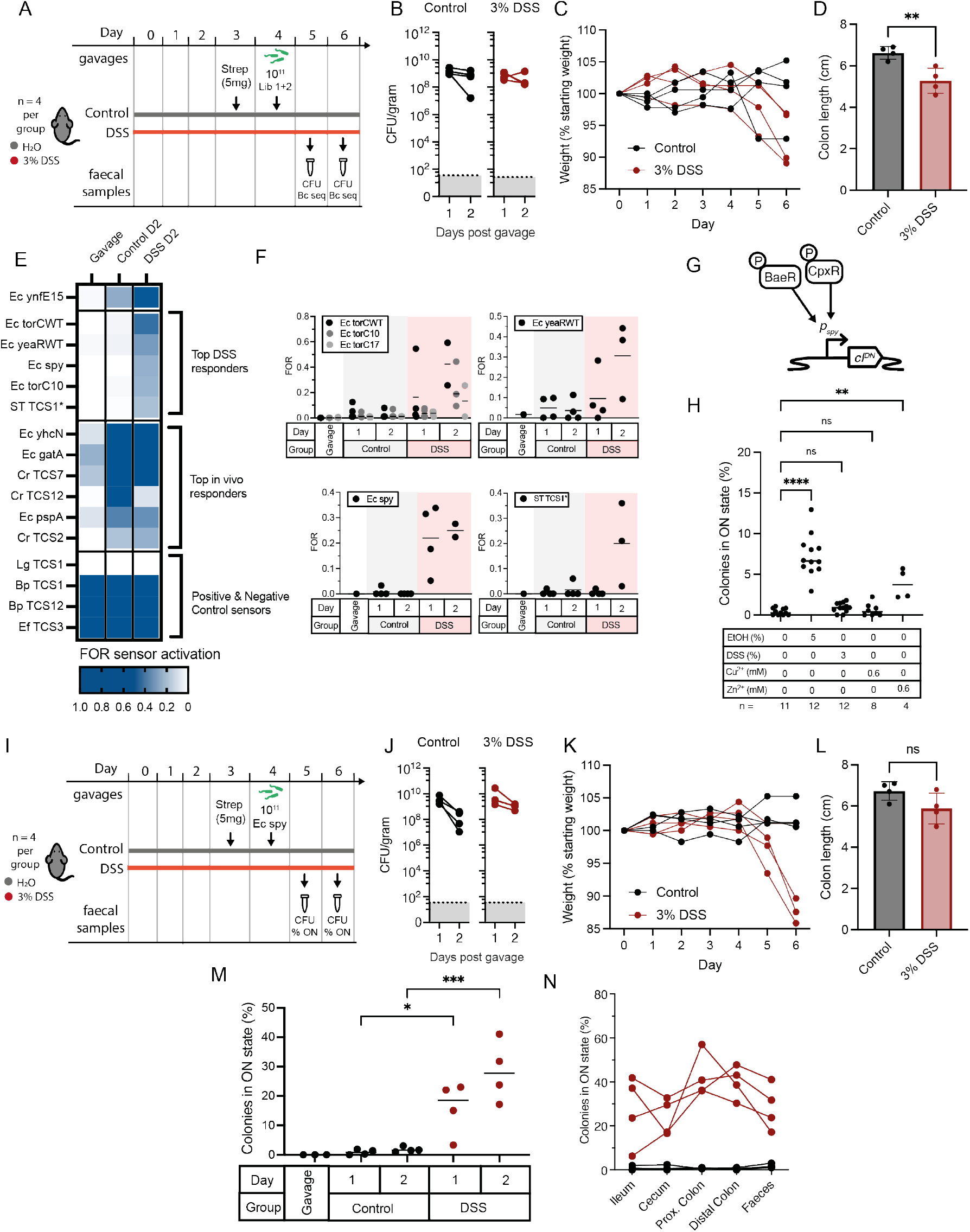
Identification and validation of inflammation-specific biosensors through screening of Library 1 + 2. A) Experimental timeline for biosensor library screening of combined Library 1 + 2 sensors in animals with or without intestinal inflammation induced by DSS administration. B) Colonisation levels in all mice were comparable. C) Fractional weight change of mice and D) colon lengths measured at the experimental endpoint on day 6 post DSS administration/ day 2 post bacterial administration were typical of DSS induced inflammation. Graph shows mean ± standard deviation and results from an unpaired T-test, p-value: 0.0075. E) Sensor activation of the top-ranked DSS responsive, in vivo responsive, and control sensors as averaged FOR across all mice on day 2 post library administration. The full dataset is shown in Supplementary Figure 6. F) individual sensor activation (FOR) from each animal for top-ranked inflammation responsive sensors. Graphs also show mean. G) Predicted regulation of E. coli P_spy_. H) Response of DTR306 Ec spy sensor when exposed to different inducers in vitro. Graphs show mean. Statistics calculated by one-way ANOVA between groups as shown, with Dunnet’s multiple comparisons adjustment. Adjusted p-values: ** = 0.001, **** = <0.0001, NS = not significant I) Experimental timeline to test DTR306 Ec spy sensor in mice with or without intestinal inflammation induced by 3% DSS administration. J) Colonisation levels between groups and animals remained high across the days following sensor bacteria administration. K) Fractional weight and L) colon lengths on day 6 post DSS administration/ day 2 post bacterial administration were indicative of DSS-induced inflammation (P value: 0.0988 by unpaired t-test). M) Response of DTR306 Ec spy sensor by plating of faecal contents on days 1 and 2 post bacterial administration showed elevated response in DSS-treated animals. Graphs show mean. Statistics calculated by one-way ANOVA between groups as shown, with Šídák’s multiple comparison adjustment. Adjusted p-values: * = 0.0195, *** = 0.0003. N) Plating of contents from different intestinal regions at experimental endpoint on day 2 post bacterial administration confirms elevated sensor memory response in all tested regions within the DSS-treated gut.

To account for the ∼10-fold increase in barcode abundance in the combined library compared with Library 1 alone (Supplementary Figure 2B-C, Data File 1), increased volumes of faecal pellet supernatants (corresponding to 30 – 40 mg worth of faecal pellet per mouse, and approximately 10^7^ CFU of engineered bacteria as estimated from colonisation levels) were inoculated into outgrowths before splitting into +/-spectinomycin cultures. Following barcode sequencing, fractional odds ratios were then calculated at the sensor level to account for the expected stochasticity of the lower abundance barcodes within the recovered fractions.

Several sensors showed elevated FOR in response to either the mouse gut or the DSS-treated gut specifically (Figure 4E, full dataset: Supplementary Figure 6A). The top 5 DSS responsive sensors included two *E. coli torC* promoter variants (with wt or synthetic 5’UTR MCD10), likely activated by trimethylamine-N-oxide ^26^; *E. coli yeaR* promoter, potentially activated by nitrate, nitrite or nitric oxide ^27^; *E. coli spy* promoter, reported to be regulated by BaeR and CpxR ^28,29^ and previously identified as being upregulated in response to DSS treatment ^8^; and, ST TCS1* (TtrRS-P_ttrBCA_) (see Figure 2), previously implicated in tetrathionate sensing during inflammation ^6^ (Figure 4E-F). As expected, the control strain, Ec *ynfE15*, showed activation in all animals, with stronger sensor activation in faecal samples from day 2 (Figure 4E, Supplementary Figure 6A-B). Confirming our previous results (Figure 3), top-ranked sensors from Library 1 that activated in all animals included Cr TCS7 (QseCB-P_ygiW_) and Cr TCS2 (DpiAB-P_*citC*_). Additional gut-responsive sensors of interest identified in this screen include the promoter region of *E. coli yhcN*, reported to be induced by cytoplasmic acid response ^30^, Cr TCS12 (*C. rodentium* NarXL TCS and diverging promoter region to the NarK homologue), and the promoter regions of *E. coli gatA* and *pspA* genes (Figure 4E, Supplementary Figure 6A-B).

Of the top-ranked inflammation responsive sensors, Ec *spy*, showed particularly low response in control animals and consistent response across DSS-treated animals (Figure 4F). To explore its induction behaviour in the context of our memory circuit more extensively, we therefore cloned and tested the sensor individually (DTR306). DTR306 cultures were grown for 4 hours in aerobic SOC broth in the presence of previously reported inducers of the *spy* promoter: EtOH (5%), zinc (0.6mM) and copper (0.6mM) (Figure 4H) ^28,31^. Colony screening on indicator plates demonstrated clear induction of DTR306 by ethanol and zinc, but not copper. Neither did we see *in vitro* induction of DTR306 in the presence of 3% DSS, indicating the substance itself is not inducing sensor activation (Figure 4H).

To confirm *in vivo* sensor behaviour, DTR306 was administered by oral gavage as an individual strain to mice. Animals (n = 4 per group) were treated with or without 3% DSS in drinking water for four days (∼10^11^ bacteria/ mouse) (Figure 4I), and streptomycin was provided (5mg/ mouse by oral gavage) 24 hours prior to DTR306 administration. Colonisation levels were high across the 2 days post administration in both groups (Figure 4J). Weight loss throughout the experiment was indicative of the impact of DSS treatment (Figure 4K). Average colon length was also reduced in DSS-treated mice, albeit without statistical significance (Figure 4L). Analysis of DTR306 response via growth on indicator plates demonstrated consistent increase in switching specifically in DSS-treated mice on both days (Figure 4M), consistent with the library -based screening results. Growth of DTR306 from endpoint dissection of the gut also demonstrated consistent switching in all regions of the gut (Figure 4N). Together, these results demonstrate the potential for the library-based construction and screening approach to identify compatible inducers for the development of memory biosensors responsive to a multitude of conditions.

## Discussion

Our work expands the use of a robust bacterial memory circuit to simultaneously evaluate hundreds of uniquely barcoded biosensor strains in the murine gut or *in vitro* culture. By incorporating a DNA barcoding strategy, using a single, hyper-variable oligonucleotide for barcode delivery, we generate large libraries of diversly sized sensors in a low-cost, single-pot reaction (Figure 1). Barcoding also facilitates iterative addition, combination and revision of libraries over time, as demonstrated throughout this current project. The successful identification and validation of several inducible circuits, both novel and previously identified, highlights the effectiveness of our pipeline in facilitating biotechnology tool development (Figure 2-4).

Novel sensor development is an area of need for synthetic biology in the quest to expand the toolbox of health- and disease-relevant inducible circuit components. Barcoded HTMS analysis is compatible with any transcriptional activation-based sensing circuits that function in the commensal *E. coli* host. This is a particular strength of our method, sourcing and testing diverse heterologous sensing components to drive memory circuit activity. By comparison, another powerful parallelised bacterial memory approach, RecordSeq ^8,32^, provides a transcriptome-wide view of chassis cell gene expression, akin to a recording of an RNAseq dataset over time. The flexibility of GGA for allowing simultaneous assembly of multiple fragments, facilitates future expansion of the approach to more complicated sensor designs, for example non-grouped TCSs, or those that regulate non-proximal or multiple promoters throughout the genome. Furthermore, use of alternative type IIS restriction enzymes can be used to expand cloning compatibility of sensors, as demonstrated here during Library 2 construction which used both SapI and BsaI enzymes.

The cloning strategy afforded libraries of thousands of uniquely barcoded strains, encoding hundreds of variant sensors. Not all targeted sensors were successfully cloned, with lack of PCR amplification the greatest source of loss within the pipeline. This is indicative of the challenges of undertaking PCR primer design and reactions in bulk using diverse bacterial genomes. Of the amplified sensors, >80% were successfully assigned to at least 1 barcode in the library. A fraction of the sensors were constitutively ON in all tests, which is expected in a memory system for any gene promoters that are routinely activated during *in vitro* culture conditions. In some cases, this may also result from a lack of native regulation and thus high basal expression of cI^DN^ in the heterologous host. TCSs derived from species more closely related to the *E. coli* NGF-1 host were, unsurprisingly, more likely to show signs of functionality, consistent with systematic assessment of porting non-native genetic systems into new hosts^33^. Many promoters are controlled by combinations of TCSs, and most TCSs regulate gene expression from a variety of targets ^34^, suggesting that cross-regulation between host and heterologous TCSs and gene expression is both likely and necessary. Tetrathionate sensing, driven by TtrRS regulation of the adjacent *ttrBCA* promoter from *S*. Typhimurium ^6^ or *E. tarda* (Figure 2F) requires cross-regulation by host *E. coli* FNR. Our experiments with truncated *Cr* TCS2 sensor variants also demonstrated cross-regulation by host *E. coli* DpiAB, NarL and cAMP:CRP (Supplementary Figure 5C). Future libraries may take this systems biology perspective into account by focussing on candidate TCSs from closely related species. Together, these challenges point to the power of library screening approaches for rapid sensor identification and validation within specific engineering biology contexts.

During *in vivo* library screening experiments, population bottlenecking caused barcode and strain diversity loss following administration to the gut, ultimately limiting experiment length (Supplementary Figure 3). Barcoded *C. rodentium* libraries have showed similar dynamics ^21,22^, indicating that bottlenecking is likely an inevitable physiological constraint. Nevertheless, our colonisation data suggests that library screening approaches are feasible and valuable over short experimental periods (eg. up to 2 days) to simultaneously test thousands of uniquely barcoded strains within the gut. Testing directly in the *λ*-memory system encoded in mouse commensal *E. coli* NGF-1, which has previously demonstrated low cellular burden, long-term stability, and function over many months in the murine gut ^5,6,11^ allows for rapid progression from screening to follow up tests as a single sensor in animals where these bottlenecking issues are no longer a concern (eg. Figure 2-4).

This pipeline not only enables the construction of biosensors tailored to the gut environment but also offers the potential for screening biosensors in other microbial growth permissible and potentially inaccessible environments. The ability to build and test new whole-cell biosensors at high throughput is a powerful method for the identification of novel inducible synthetic circuit components and prototype disease diagnostics.

## Methods

### Media and culture conditions

Bacterial cultures of individual strains were grown at 37°C in LB broth or on LB agar plates. Library cultures were grown in LB peptone special (LBPS) containing 10 g/L peptone (Sigma product no. 68971), 10 g/L NaCl, and 5 g/L yeast extract in deionized water. LBPS was used because it does not contain lactose so avoids the potential for biased outgrowth of ON sensors expressing *lacZ*. For anaerobic growth, cultures were grown statically in pre-reduced media in a BD GasPak EZ container system with container system sachets to generate the anaerobic environment (BD, product no. 260001) or using an InvivO_2_ workstation with I-CO_2_N_2_IC gas controller fitted for anoxic conditions (Baker Ruskinn). For memory response quantification and blue-white screening, cultures were plated on LB agar plates containing 60 μg/mL 5-bromo-4-chloro-3-indolyl-β-d-galactopyranoside (X-gal; Thermo Fisher Scientific catalogue no. R0402). Antibiotics in media for bacterial growth were used at the following working concentrations; streptomycin (Sigma-Aldrich, S650) 200 μg/mL, chloramphenicol (Sigma-Aldrich, C0378) 25 μg/mL, carbenicillin (ThermoFisher Scientific, BP2648) 100 μg/mL, and spectinomycin (MP Biomedical, 158993) 50 μg/mL.

Individual biosensor inductions *in vitro* were conducted in SOC media (2% Tryptone, 0.5% yeast extract, 10 mM NaCl, 2.5 mM KCl, 10 mM MgCl_2_, 10 mM MgSO_4_, 20 mM glucose) with the specified concentration of inducer: potassium tetrathionate (Sigma-Aldrich, P2926), sodium citrate tribasic dihydrate (Sigma-Aldrich, 71406), sodium nitrate (VWR, 27955), copper chloride dihydrate (Sigma-Aldrich, C6641), or zinc sulphate (Sigma-Aldrich, Z0251). SOC media was used for individual sensor testing for comparability with previous individual memory sensor testing (Naydich 2019, Riglar 2017). M9 (1x Difco M9 Minimal Salts (Scientific Laboratory Supplies,: 248510), 0.1 mM CaCl_2_, 1 mM MgSO_4_, 0.4% filter sterilised glucose, 0.2% casamino acids (Fisher Scientific Fisher,12747109) and 1μg/mL niacinamide) media was used for *in vitro* library screening.

### Computational pipeline for TCS sequence extraction

A custom Python script was used to extract TCS DNA sequences from the Microbial Signal Transduction Database (MiST 3.0) ^35^ and NCBI RefSeq databases. All HK and RR TCS signalling genes were extracted from MistDB for a given bacterial strain and pairs of partner HK and RR were identified by proximity (<200bp apart) and orientation (both genes must be on the same strand). Genes neighbouring the paired TCS were considered likely regulated targets if they were encoded adjacent to, and on the opposing strand of, the HK and RR pair. The strain’s DNA sequence and annotations were automatically downloaded from Refseq’s FTP site. The TCS’s gene IDs from MistDB were then matched to the RefSeq annotations and the combined DNA sequence for the TCS and intergenic region between the TCS and its putative target were extracted. Next, the combined TCS & Target sequences were screened for SapI or BsaI sites, and, if absent, were deemed ‘SapI or BsaI compatible’. Primers for compatible TCSs were designed using Primer3 (in Geneious Prime software version 2020.1.1-2023.1.1; Biomatters Ltd) with SapI or BsaI overhangs for Golden Gate Assembly (GGA). A full list of predicted grouped TCS amplicons (including HK, RR and intergenic region) and primers used for TCS amplification is provided (Data File 1).

### TCS Library construction

Additional oligonucleotide and primer sequences used for library construction and sequencing (Table 2) and bacterial strains used throughout the study (Supplementary Table 3) are provided.

**Table 2:**
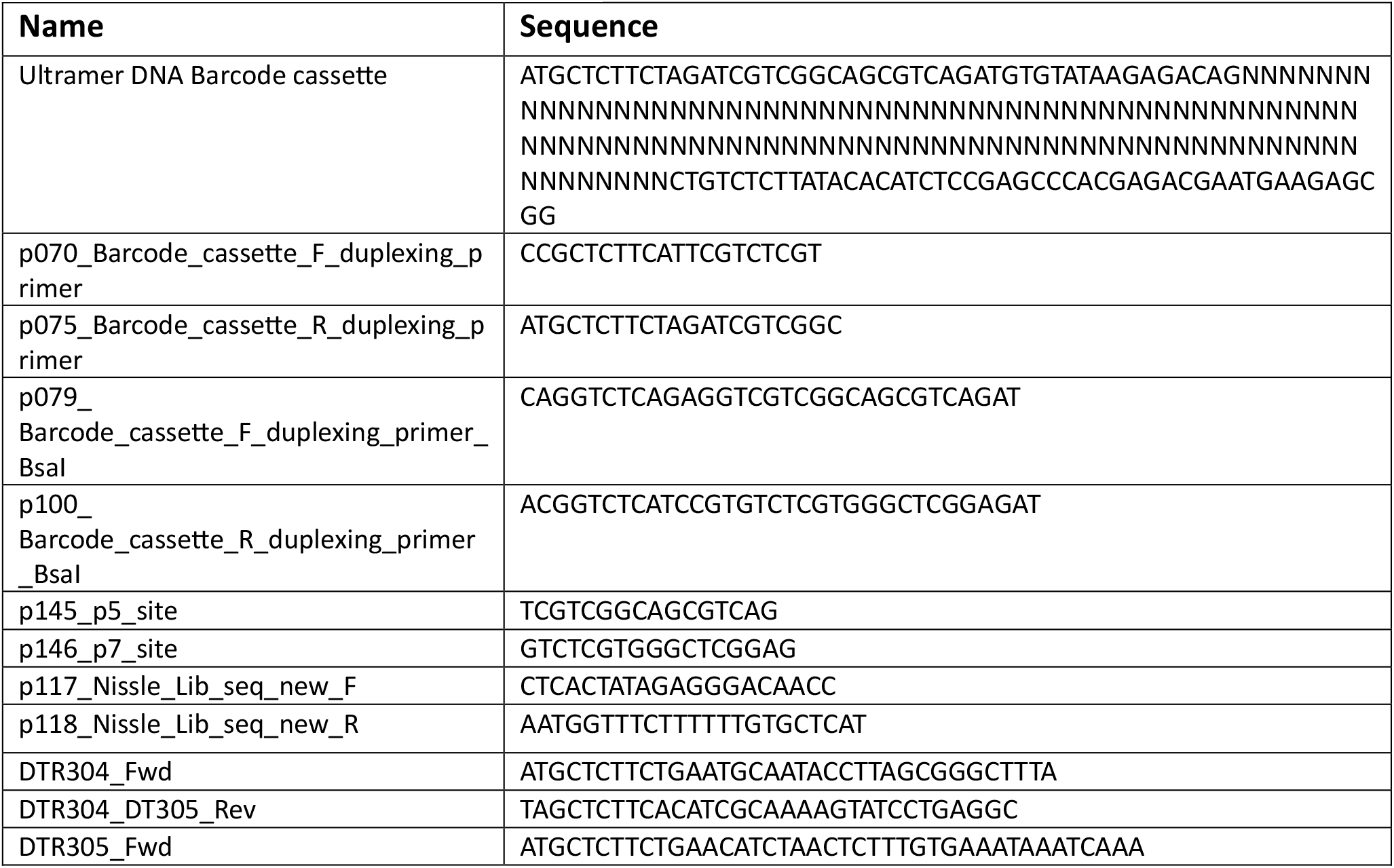
Primers and DNA fragments.

**Table 3:**
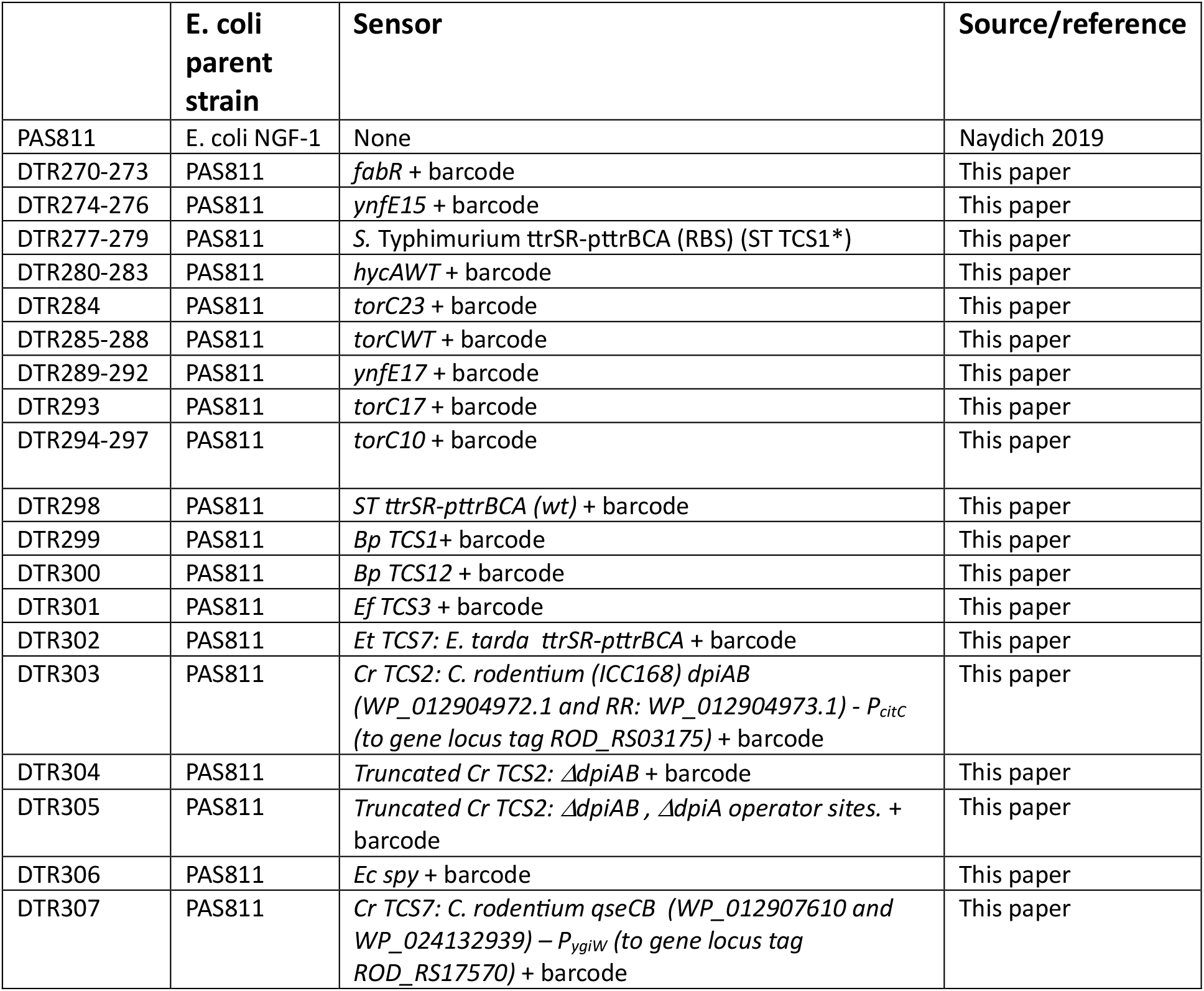
Bacterial strains.

Purified bacterial gDNA was sourced from Public Health England (NCTC 10396 *Edwardsiella tarda* and NCTC 11829 *Blautia producta*), by donation from other laboratories, or, for *Citrobacter rodentium*, purified using the Gentra Puregene Yeast/Bacteria gDNA extraction kit (Qiagen catalogue no. 158567). TCSs were amplified by PCR using Q5 High-Fidelity DNA polymerase (New England Biolabs), and amplified PCR fragments were purified by column purification using the Monarch PCR Cleanup Kit (New England Biolabs) following the manufacturer’s protocols.

DNA barcodes were synthesised as an Ultramer DNA Oligo (IDT) consisting of a 106bp variable nucleotide region flanked by Illumina sequencing primer binding sites, allowing direct sequencing of the barcode region, and SapI restriction sites, for GGA assembly (Table 2). The DNA barcode oligo was made double stranded by Q5 PCR with 5 cycles with 8 pmol of starting oligo template and 32 pmol of each duplexing primer (primers p070 and p075 for SapI based GGA, and p079 and p100 for BsaI based GGA, in Table 2).

The destination vector was pCR06 (for SapI assembly), a modified Tn7 transposon backbone derived from pGRG36 ^36^ via pDR07 ^11^, or pDR07 (for BsaI assembly). Amplified sensors were pooled and assembled with the double stranded barcode oligo into the Tn7 transposon machinery containing backbone (pCR06 or pDR07) by multiplexed GGA using SapI or BsaI. SapI GGA reactions were set up using: 3nM of each DNA fragment, 3μL(15 units) of SapI, 0.25μL(500 units) of T4 DNA Ligase, 2μL of T4 DNA Ligase buffer and H20 to a final volume of 20μL. BsaI GGA reactions were set up using: 3nM of each DNA fragment, 1.5μL (30 units) of BsaI-HFv2, 0.5μL (1000 units) of T4 DNA Ligase, 2.5μL of T4 DNA Ligase buffer and H_2_0 to a final volume of 25μL. Cycling conditions for all GGA were: (37°C, 5 min and 16°C, 5 min) x 30, followed by 60°C, 5 min, then storage at 4°C. For each library, 4 μL of the GGA mix was transformed into 50 μL of ElectroMAX DH5α-E competent cells (Thermo Fisher Scientific) by electroporation according to the supplier’s protocol and plated onto 5 carbenicillin selective LB-agar plates. An additional 1:100 dilution of the transformation reaction was plated to estimate library size. All transformants were scraped from LB-agar transformant plates using 1 mL of pre-warmed LB, vortexed to mix thoroughly, and frozen as a glycerol stock. A large inoculum of the glycerol stock was grown in 5 mL of LB overnight, miniprepped and transformed into electrocompetent E. coli NGF-1 encoding HTMS memory (PAS811). Transformant plates were scraped with pre-warmed LB, pooled, diluted 1:1000 and grown for 6 hours at 30°C in LBPS-streptomycin-carbenicillin-chloramphenicol with 0.1% L-arabinose to induce Tn7-based genome integration. Cells were then diluted 1:1000 and grown overnight at 42°C three times for plasmid curing. Libraries were frozen as 25% glycerol stocks until use.

Individually cloned sensors (Ec yeaRWT, Ec yeaR23, Ec ynfE17, Ec torC10, Ec torC17, Ec torC23, Ec torCWT, Ec hycAWT), control sensors (Ec ynfE15 and ST TCS1*), and fabR positive normalisation strains were cloned using identical protocols to the library construction, with the exception of using individual sensor PCRs for GGA reactions (Data File 1) and picking multiple separate (so separately barcoded) individual colonies for plasmid integration and curing steps. Sensors and barcodes of individually cloned constructs were verified and identified by sanger sequencing (Eurofins) (Data File 1). Correctly assembled individually cloned sensors were combined with the pooled library at ∼1:1000.

### Truncation mutant cloning

DpiA binding sites within the Cr TCS2 intergenic promoter region were identified by sequence alignment to the corresponding genomic region in *E. coli* MG1655 within the Ecocyc database ^17^. Plasmids for truncated sensor mutants DTR304 and DTR305 were cloned by PCR amplifying appropriate regions (using DTR304_Fwd and DTR304_5_Rev or DTR305_Fwd and DTR304_5_Rev respectively), assembled by GGA using SapI along with unique barcodes, and transformed and genome integrated into PAS811 as above.

### Library sequencing and barcode assignment

For nanopore long read sequencing, the barcode and sensor regions of the biosensor libraries were amplified using Q5 PCR (New England Biolabs) directly off 5 μL of glycerol stock using primers p117 and p118 (Table 2). PCR products were purified using a Monarch PCR Cleanup Kit (New England Biolabs) and prepared for nanopore long read sequencing using the Ligation Sequencing Kit (Oxford Nanopore Technologies, SQK-LSK109). Library 1 was run on a Flongle flow cell (R9.4.1) and Library 2 was run on a Minion flow cell (R10.4.1). Basecalling from Fast5 files was completed using MinKNOW software (version 22.12.5). Nanopore adapters were trimmed in Geneious (adapter sequence: CCTGTACTTCGTTCAGTTACGTATTGCT) with 5bp allowed mismatches, minimum 10bp match to adapter and 0.1 error probability limit (regions with more than a 10% chance of error per base were trimmed).

Trimmed nanopore data was filtered for reads >400bp (a read length sufficient to at minimum partially cover both the barcode and sensor regions), aligned to TCS reference sequences (excluding barcode regions) of all successfully PCR amplified TCS using Minimap2 (version 2.17). Long read basepairs that extend past the TCS reference sequence were trimmed using a custom python script and aligned to the inferred barcodes from short read sequencing data (see below) using Bowtie2 ^37^. Barcodes were assigned to a specific sensor (Data File 1) if that sensor had both the highest number of trimmed long reads aligning to the barcode, and highest fraction of aligning trimmed long reads normalised by total long reads for the specific sensor, with a minimum of 5 long reads for assignment. Any barcodes with different assignments between those values, or to which no trimmed long read data aligned were discarded for odds ratio calculations.

For short read sequencing, genomic DNA (gDNA) was extracted from ± spectinomycin overnight cultures manually using the Gentra Puregene Yeast/Bacteria gDNA extraction kit (Qiagen catalogue no. 158567) or on a FeliX liquid handler using the innuPREP DNA/RNA kit with an initial manual lysozyme step for bacterial cell lysis. For the lysozyme bacterial cell lysis steps, 500μL of bacterial overnight culture was pelleted, resuspended in 200μL of enzymatic buffer consisting of 20mM TrisCl pH 8, 2mM sodium EDTA, 1.2% Triton X-100, and 20mg/mL lysozyme and incubated at 37°C for 30 minutes. Barcodes were first Q5 PCR amplified from extracted gDNA using primers p145 and p146 (Table 2), then purified by AMPure XP beads at a ratio of 1.8x beads to sample following the manufacturer’s protocol (Beckman Coulter). Purified barcodes were then Q5 PCR amplified (using a modified protocol with 12 cycles) using IDT UDIs and purified again by AMPure XP beads, as above. Final fragments for short read sequencing were quantified using Qubit dsDNA BR assay kit (Invitrogen) following the manufacturer’s protocol and pooled at equimolar concentrations. Sequencing was undertaken on an Illumina NovaSeq sequencer (150bp Paired-end) (Novogene), with an average of 1.4 million reads per sample.

### Short read sequence barcode analysis

Short read sequencing of library screening experiments was analysed using Dada2 ^38^ with a custom R script to generate inferred barcodes and barcode counts for each sample (settings: truncLen=106, max expected errors maxEE = 1, truncQ=2). Inferred barcodes were filtered to the expected barcode length of 106bp and clustered on 95% sequence similarity using the seq_cluster function from bioseq ^39^. The exact barcode sequence with the highest read number in each cluster was kept as the cluster’s reference barcode and read counts from other barcode members of the cluster added to the reference barcode.

### Fractional Odds ratio calculation and sample QC

Sensor activation is calculated using a fractional odds ratios defined for each sensor. First, the odds ratio for each barcode in a sample is calculated as:

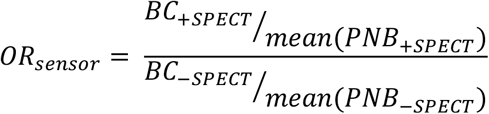

Where BC_+SPECT_ and BC_-SPECT_ are reads from an individual barcode from samples grown with and without spectinomycin respectively, and PNB_+SPECT_ and PNB_-SPECT_ are combined read numbers from several positive normalisation barcodes in samples grown with and without spectinomycin respectively. Odds ratios were initially calculated using the averaged results from four barcoded strains triggered by our previously used 100% positive normalisation promoter from *E. coli* MG1655, pFabR ^11^. However, in certain samples we observed variability in the outgrowth of spiked-in strains compared to sensors contained within the library, which appeared to grow more slowly (Supplementary Figure 7A). We hypothesised this may result from the differential growth conditions of normalisation strains and library prior to mixing, leading to differential lag phases prior to growth. Thus, we developed a more robust normalisation strategy using 14 barcodes linked to 3 sensors contained within the library (Bp TCS1, Bp TCS12 and Ef TCS3) that consistently showed odds ratios ∼1 when normalised using fabR strains, and which were subsequently validated as constitutively switching ON by independent cloning and indicator plating (Supplementary Figure 7C). When samples were normalised using these barcodes instead of the fabR barcoded strains it corrected skewness that could lead to shifted odds ratio (Supplementary Figure 7B).

Because spectinomycin is bacteriostatic, reads from OFF sensors may remain in the culture at an abundance determined by the fold change of the population’s outgrowth. This will vary based on the dilution factor used (typically 1:1000 for *in vitro* data, 1:100 for *in vivo* data) and *E. coli* cell density in the original sample (which is variable). Therefore, we further calculated a fractional OR (FOR) that takes into account observed differences in outgrowth, defined as:

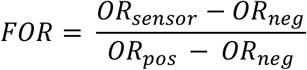

Where OR_pos_ is the average odds ratios of all positive control strains (Ef TCS3, Bp TCS12 and Bp TCS1), and OR_neg_ is the average odds ratio of an internally selected negative control (OFF) strain. Lg TCS1 was selected as an internal negative control, as it was the sensor with >5 assigned barcodes that consistently showed the lowest odds ratios across all analysed samples.

Finally, a set of QC parameters was developed to ensure any given sample would provide reliable sensor activation quantification. Replicate barcodes of individual sensors within the library allowed assessment of odds ratio variability caused by differences in bacterial starting concentration. Unsurprisingly, low concentration samples display higher variability in their outgrowth, resulting in higher deviation from the expected odds ratio of 1 for the 14 positive barcodes (Supplementary Figure 8G, H, I). Slope and R^2^ of a linear fit of OR of positive control strains are calculated to exclude samples with high levels of observed outgrowth variation (thresholds set at an R^2^ >0.6 and slope>0.5 and <2). We additionally set a conservative rationally calculated sensitivity threshold. The libraries contain sensors with abundances across ranges of orders of magnitude. We filtered barcodes to be above a set threshold of an abundance in the final sequenced -spectinomycin sample of >1/5000 of the fraction of reads. This is because any fractions below this threshold would indicate tens to a couple of hundred bacteria’ cells being split between +spectinomycin and -spectinomycin cultures (from a stationary phase *E. coli* culture containing 10^8^ – 10^9^ CFU/mL, a sensor at 1/5000 of the library is at a concentration of around 2*10^4^ – 2*10^5^ CFU/mL, after a 1:1000 dilution into ± spectinomycin cultures would be at 20 – 200 CFU/mL). To avoid any impacts from low abundance positive normalisation barcodes, barcodes with <100 reads in ± spectinomycin cultures from a given sample were excluded.

Due to the roughly 10-fold higher number of barcodes in Library 2, reads from all barcodes for a specific sensor in these samples were pooled before OR, FOR and QC calculations.

### High throughput in vitro library screening with Tetrathionate

Approximately 15μL of a glycerol stock of Library 1 was inoculated into a 4mL LBPS overnight culture with streptomycin, then back diluted 1:1000 into the relevant test condition in 0.5mL of LBPS in a 96-well deep-well plate and grown for 4 hours at 37°C aerobically or anaerobically. For aerobic growth plates were shaken at 1200 RPM. Liquid cultures of barcoded fabR strains for positive normalisation were spiked in at 1:100 and cultures from deep well plates were then split and back diluted 1:100 into 0.5mL cultures ± spectinomycin and grown overnight aerobically. Cells were pelleted and gDNA extracted using the FeliX liquid handling robot (as above).

### In vivo library and individual strain screening

All animal experiments were carried out under approval of the local Ethical Review Committee at Imperial College London according to UK Home Office guidelines. Experiments were conducted using 7-9 week old C57BL/6J female mice (Charles River UK). Upon arrival at the facility mice were given one week to acclimatise to housing conditions and diet. All animals were sacrificed humanely by cervical dislocation at the endpoint of the experiments. Mice were provided with Teklad 2019 Global Rodent Diet (Envigo). For all experiments, 24 hours prior to engineered bacterial administration, mice were administered 5mg, 1mg or 0mg of streptomycin sulphate as specified (VWR, 0382-EU-50G) by oral gavage in 100μL PBS and confirmed free of streptomycin resistant bacteria by faecal plating on streptomycin selective LB agar plates 24 hours post streptomycin oral gavage but prior to engineered bacterial delivery. Where specified, colitis was induced via administration of 3% dextran sodium sulphate in drinking water for the specified number of days before bacterial gavage.

For library screening experiments, libraries were grown overnight, washed in PBS and concentrated 10x (Library 1) or 5x (Library 1 + 2). 200μL of the washed library were then administered by oral gavage (∼10^10-12^ cells per mouse). For sensor activation and bacterial load quantification, fresh faecal pellets were collected and stored immediately in 100μL of PBS, then resuspended to 100 mg/mL in PBS, vortexed for 2 mins, pulse centrifuged to remove large debris, before being diluted and plated to track bacterial load. The remainder of the faecal supernatant (up to a limit of 400 μL for Library 1) was inoculated into 4 mL streptomycin chloramphenicol LBPS and grown for 4 hours. For Library 1, positive normalisation fabR strains were also diluted 1:1000 from overnight cultures and grown for 4 hours, before being diluted 1:100 into 5mL of the faecal pellet outgrowths. After thorough mixing, cultures were diluted 1:100 into 10mL fresh LBPS, split into 2x 5mL culture, to which spectinomycin was added to one, and grown overnight. Overnight cultures were pelleted, and gDNA extracted and amplified for short read sequencing.

For testing Ec spy, mice were administered 3% DSS for the specified number of days before bacterial gavage. To test sensor activation, faecal pellets were collected in 100 μL of sterile PBS, then resuspended to 100 mg/mL, vortexed, pulse centrifuged to remove large debris and diluted and plated on indicator X-gal LB-agar plates for blue-white screening. To test sensor activation within different regions of the gut, at the endpoint of the experiment mice were dissected and the entire contents including mucus of the small intestine, cecum, proximal colon or distal colon were scraped, collected, resuspended to 100 mg/mL, vortexed, centrifuged, diluted and plated as for faecal pellets. The proximal colon was selected as the top half of the colon (from the bottom of the cecum to halfway to the anus), and distal colon is defined as the bottom half of this region.

### Statistics and Sensor Rankings

Statistical tests were performed in Graphpad Prism v9 or v10 software. For ranking responsive sensors in the Library 1 *in vivo* screen (Figure 3D) and Library 2 *in vivo* screen (Figure 4E), sensors were first filtered for those OFF in the gavage sample (FOR <0.25). To rank *in vivo* responsive For Library 1 screening (Figure 3), sensors were ranked by greatest difference between average faecal FOR across all mouse samples and gavage FOR. Sensors were then classified as ‘top *in vivo* responders’ if they had a response higher than the control Ec ynfE15. To rank inflammation responsive sensors in the Library 1 + 2 screen (Figure 4), sensors were ranked by difference in average faecal FOR in -treated animals (D2) and average faecal FOR in control animals (D2). Sensors were classified as ‘top DSS responders’.

## Data availability

TCS DNA sequences, including intergenic regions up to neighbouring genes and primer sequences used to amplify the TCS, and library barcode sequences are available in Supplementary Data File 1.

## Code availability

All custom code used to analyse sequencing data or search relevant databases and extract TCS are publicly available at https://github.com/ClareRobin/Robinson2023_BiosensorsLibrary.

## Acknowledgements

We would like to thank R Jackson, M Crone, N Matthews, and K Rekopoulou for assistance with sequencing and sequencing analysis; M Tran and K Jensen for assistance with automation; G Frankel, J Marchesi, C Mullineaux Sanders, L Roberts and D Chrystomou, for gifts of, and assistance with, genomic DNA. PAS811 was a gift from P Silver. This research was funded by a Sir Henry Dale Fellowship to D.T.R (211230/Z/18/Z). C.M.R. was funded by a President’s PhD Scholarship through Imperial College London.

## Author contributions

D.R., C.M.R and D. C. conceived of the study and experimental designs. D.R. and C.M.R. wrote the manuscript. All authors contributed to the editing of the manuscript. T.W. and D.R. aided library construction design, and C.M.R finalised and cloned all sensors. C.M.R. performed sequencing and computational analyses. C.M.R and D.C. performed the animal experiments. Y.C. built and characterised the C. rodentium DpiAB-p_citc_C variants.

