## Supplementary Figures 1-8 for "A discovery platform to identify inducible synthetic circuitry from varied microbial sources"

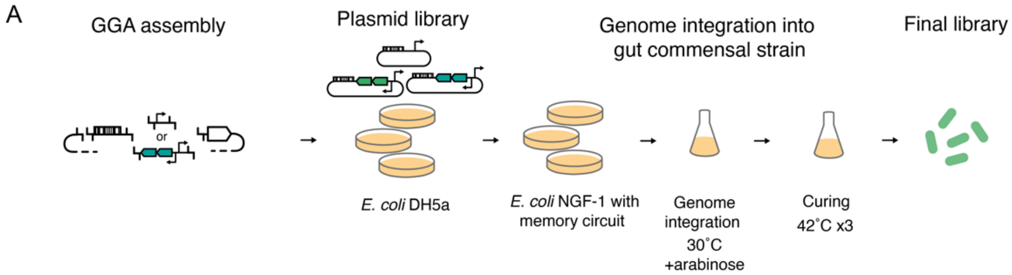


**Supplementary Figure 1 Pooled cloning strategy.** (A) Libraries were constructed by multiplexed golden gate assembly to create a barcoded plasmid library, which was first transformed into *E. coli* DH5a, then miniprepped and transformed into *E. coli* NGF-1 PAS811, a murine gut commensal *E. coli* strain carrying the high-throughput memory circuit. Induction of Tn7 genome integration machinery by arabinose and curing of plasmids by multiple growth steps at 42°C afforded the final pooled library.

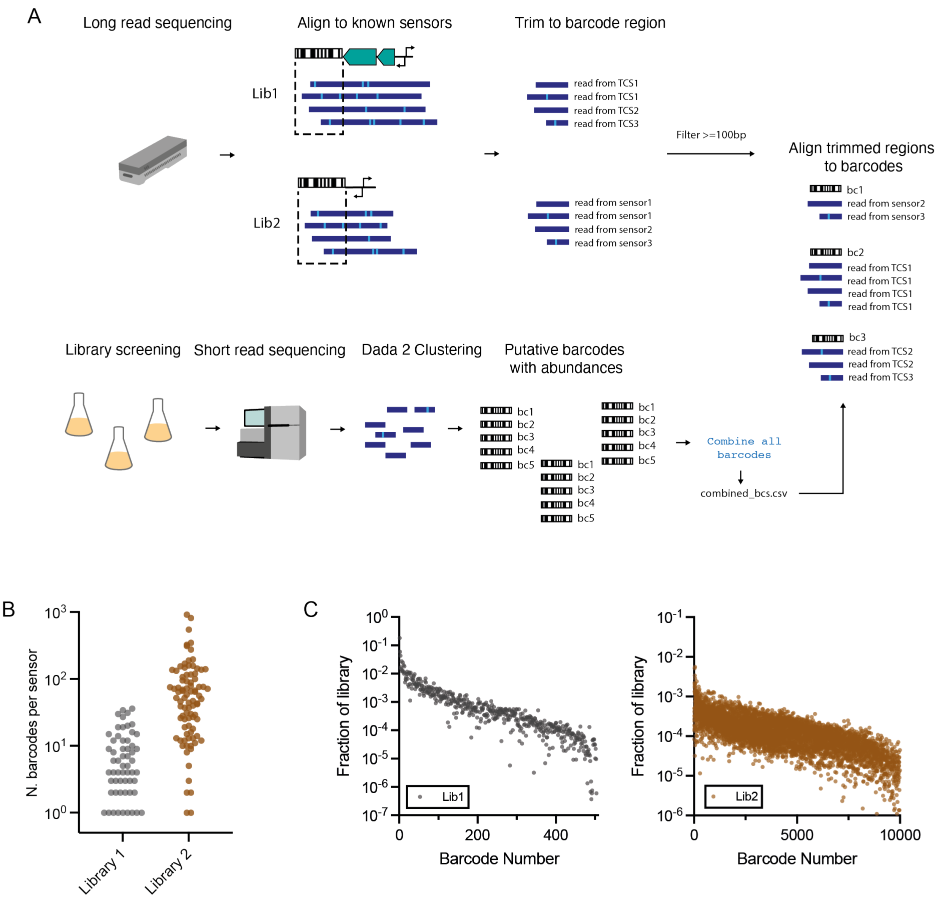


**Supplementary Figure 2: Barcode assignment.** A) Barcodes were assigned to sensors by a combination of Nanopore long-read sequencing and Illumina short-read sequencing, from which high confidence barcodes were inferred using Dada2. Long-read data were first aligned to anticipated sensor templates, barcode regions were then trimmed and aligned to short-read inferred barcodes. A barcode was assigned to a sensor if it had both the highest number of trimmed long reads and highest fraction of aligning trimmed long reads normalised by total long reads for the specific sensor. B) Number of assigned barcodes per sensor within libraries 1 and 2. C) Fractions of total reads across all samples of a given library's assigned barcodes.

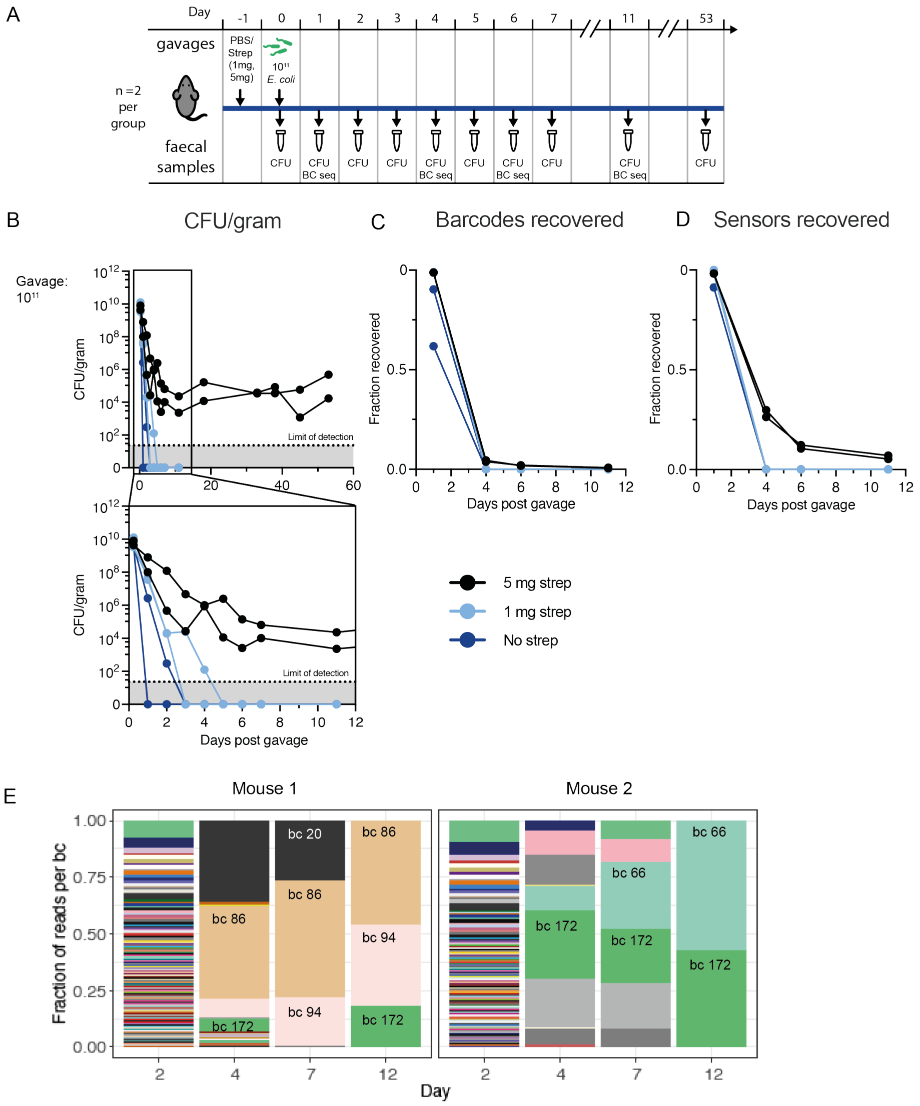


**Supplementary Figure 3. Barcode and strain diversity is maintained over short periods in the murine gut**. A) C57BL/6 mice (n=2 per group) were administered different concentrations of streptomycin (0, 1 or 5 mg) via oral gavage, followed by ~$10^{11}$ Library 1 bacteria the following day. Faecal samples were taken on days following administration for colonisation and diversity analysis, measured by plated CFU counts and Illumina barcode sequencing, respectively. B) CFU counts of engineered bacteria, C) fraction of total library barcodes and D) fraction of total sensors recovered across the experiment. E) Relative abundance of each barcode within two mice treated with 5 mg streptomycin. Individual barcodes are high abundances in later timepoints are numbered and highlighted for clarity.

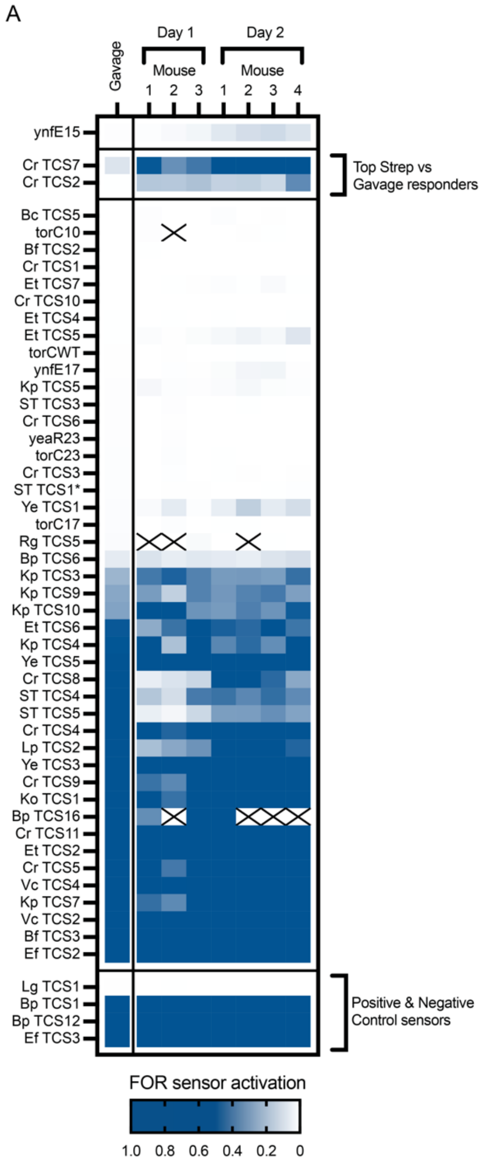


**Supplementary Figure 4: Library 1 sensor response to the murine gut environment**. A) Sensor activation (FOR) of all sensors in all mice on all days. Heatmap shows the average FOR of all barcodes for each sensor in each sample.

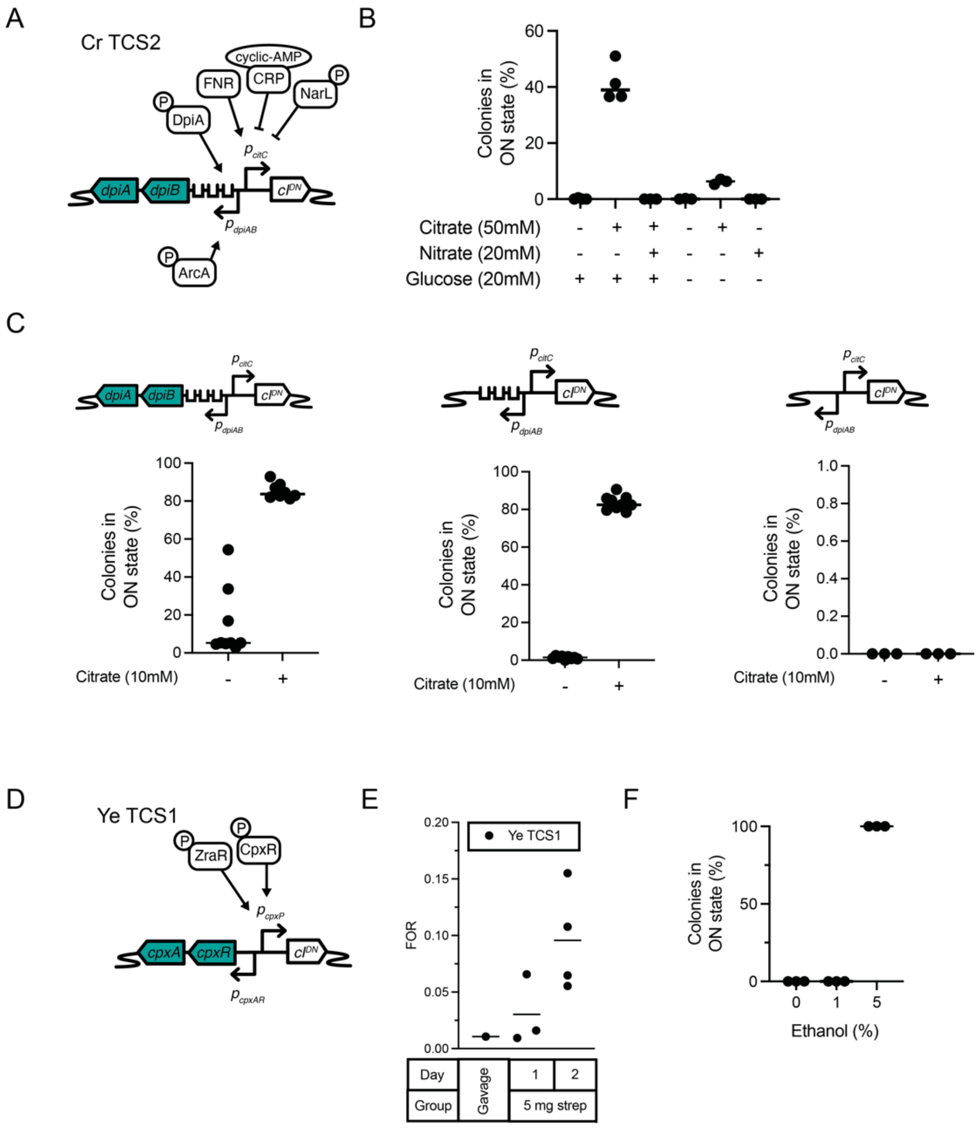


**Supplementary Figure 5: In vitro testing of in vivo identified sensors of interest.** A) Overview of predicted regulation of Cr TCS2 (DpiAB -$P_{citC}$). B) *In vitro* testing of the Cr TCS2 sensor shows it is responsive to citrate in the presence of glucose and absence of nitrate. C) Truncated versions of the sensor show that E. coli host machinery is capable of regulation, with full sensor response (left) similar to that in the absence of the heterologous *C. rodentium* derived TCS genes *dpiA* and *dpiB* (middle). However, the sensor requires DpiAB binding sites for response to citrate (right). D) Overview of predicted regulators of Ye TCS1 (CpxAR - $P_{macA}$) E) Response of Ye TCS1 in the screen of Library 1. Panel shows average FOR from gavage and faecal samples of each mouse on days 1 and 2 post administration. Mean is marked F) Testing of the sensor individually demonstrates complete activation in the presence of the previously documented inducer, 5% ethanol.

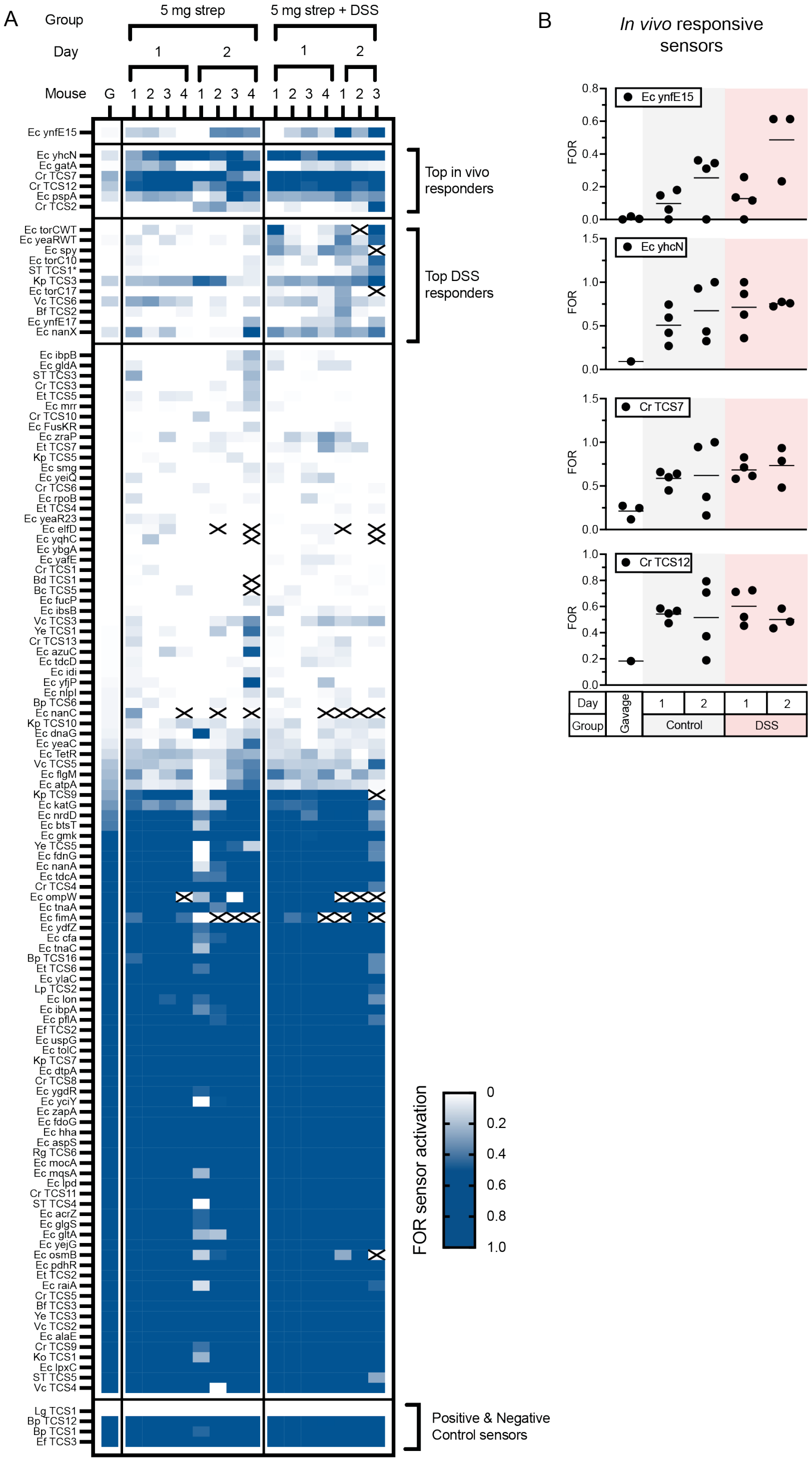


**Supplementary Figure 6: Combined Library 1 + 2 sensor response to the murine gut with and without induced inflammation** A) Full biosensor activation data for all Library 1 and 2 sensors across all mice and timepoints. Heatmap shows the FOR from pooled barcodes for each sample. B) Individual sensor activation (FOR) from each animal for the top 3 ranked in vivo responsive sensors and the Ec ynfE15 control. Graphs show mean.

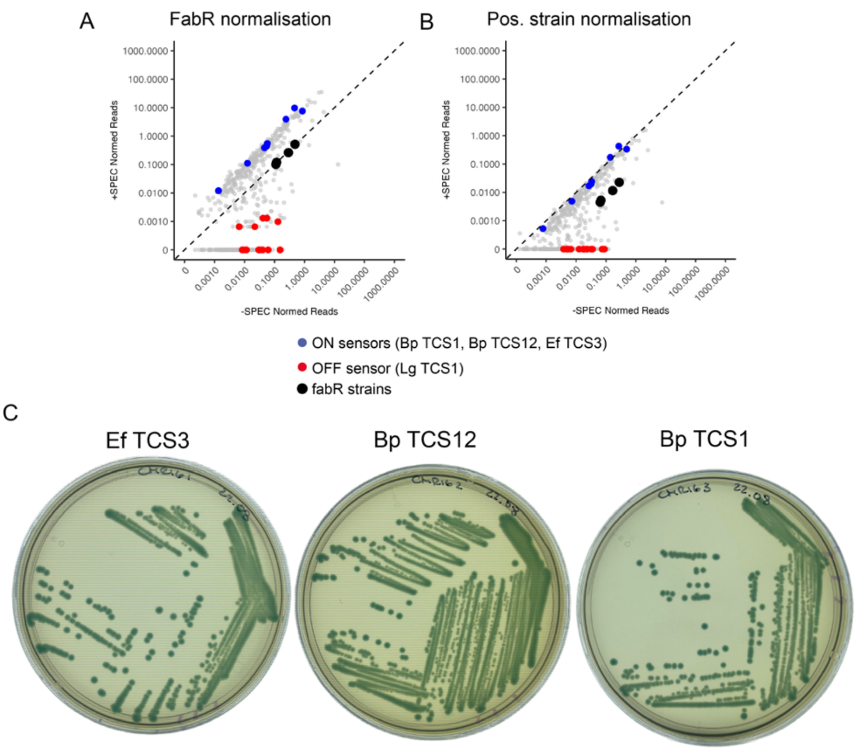


**Supplementary Figure 7: Identification of internal Positive Normalisation Strains for Odds ratio calculation** A) Example data normalised using spiked in Ec fabR controls compared to B) normalisation with internal positive control strains (Ef TCS3, Bp TCS12, and Bp TCS1). D) Individually cloned new positive control strains streaked on X-gal indicator plates demonstrated 100% memory-ON.

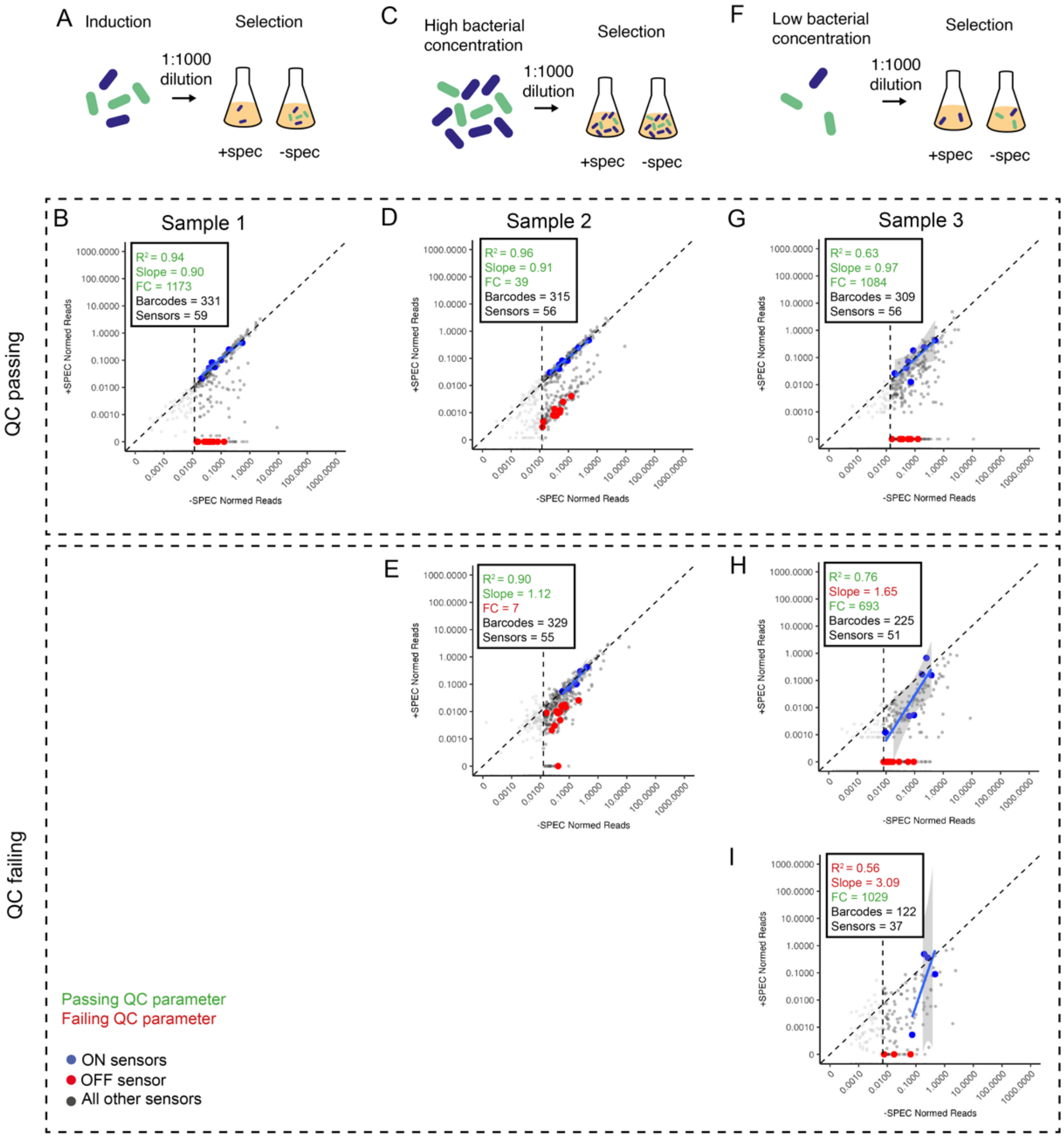


**Supplementary Figure 8: Quality Control parameters** A,B) Optimal sample conditions, in which there is a high fold-change between positive and negative strains, and positive strains show low outgrowth variability. C-E) High bacterial concentration is assumed to lead to a lack of outgrowth potential of ON strains in +SPEC samples, leading to lower fold-change between positive and negative controls strains. F-I) Low bacterial concentration at the dilution and selection stage is assumed to lead to variability in outgrowth of positive and activated strains, leading to greater variation in sensor activation distributions. QC failing samples (E), (H) and (I) have QC failing parameters marked in red.
